# Genetic Encoding of A Nonhydrolyzable Phosphotyrosine Analog in Mammalian Cells

**DOI:** 10.1101/2021.11.30.470659

**Authors:** Xinyuan He, Bin Ma, Yan Chen, Jiantao Guo, Wei Niu

## Abstract

Protein tyrosine phosphorylation plays a critical role in signal transduction and the regulation of many cellular processes. It is of great significance to understand the underlying regulatory mechanism of particular tyrosine phosphorylation events. Here we report the genetic incorporation of a phosphotyrosine (pTyr) analog, *p*-carboxymethyl-L-phenylalanine (CMF), into proteins in mammalian cells. This nonhydrolyzable pTyr analog can facilitate biological studies by removing complications caused by the dynamic interconversion between the phosphorylated and non-phosphorylated isoforms of a protein. The developed methodology was demonstrated by using the human signal transducer and activator of transcription-1 (STAT1) as a model protein for homogeneous and defined incorporation of CMF. This tool will greatly enhance one’s capability to study protein tyrosine phosphorylation-associated biomolecular and cellular events, and enhance biomedical research that target protein tyrosine phosphorylation, which will have a broad impact to both fundamental studies and practical applications.

## Introduction

Protein tyrosine phosphorylation is one of the most common and important post-translational modifications that plays a pivotal role in many aspects of cell biology. Phosphorylation of specific tyrosine residues of substrate proteins is reversibly regulated by protein tyrosine kinases (PTKs) and protein tyrosine phosphatases (PTPs). In order to study protein tyrosine phosphorylation-associated cellular events, homogeneous proteins with site-specific phosphorylations are needed. While PTKs can be used to phosphorylate proteins *in vitro*, they have limited site specificity and often results in sub-stoichiometric phosphorylation.^1^ In addition, it is challenging to identify suitable kinases for the *in vitro* modification of a large number of newly identified and putative phosphorylation sites. Semisynthetic (e.g., native chemical ligation) and cell-free methods have been developed to site-selectively introduce phosphotyrosine (pTyr).^2-6^ In addition, genetic incorporation of phosphotyrosine (pTyr) or its precursor analog in live *E. coli* cells was recently reported.^7-9^ Furthermore, synthesis of proteins containing nonhydrolyzable pTyr analogs, including *p*-carboxymethyl-L-phenylalanine (CMF) and 4-phosphomethyl-L-phenylalanine (Pmp), were also reported in *E. coli*.^7,10^ Nonhydrolyzable pTyr analogs that retain biological function but are resistant to hydrolysis by protein tyrosine phosphatases (PTPs) will greatly facilitate functional studies of tyrosine phosphorylation and associated cellular events. Indeed, the dynamic inter-conversion of phosphorylated and dephosphorylated isoforms can complicate interpretations of experimental data and further hinder deciphering the roles of individual phosphorylation sites.

A major barrier to the broad application of nonhydrolyzable pTyr analogs in biological studies is the lack of a method for their genetic encoding in live mammalian cells. Indeed, a method that enables efficient expression of proteins with defined and nonhydrolyzable tyrosine phosphorylation patterns in real-time is critical to investigations of cellular mechanisms in many pTyr-dependent signaling pathways. Here we report the first genetic incorporation of CMF, a nonhydrolyzable pTyr analog, into proteins in mammalian cells. CMF has been widely used as a reliable pTyr analog for *in vitro* studies of pTyr-associated biological processes.^11-14^ The engineered CMF-specific aminoacyl-tRNA synthetase (CMFRS) was characterized by protein expression in both yeast and mammalian cells. Its fidelity and incorporation efficiency was further studied in 293T cells. Functional replacement of pTyr by CMF was demonstrated using the human signal transducer and activator of transcription-1 (STAT1) as the model system through *in vitro* binding and immunofluorescence assays in live mammalian cells. Our research validate that CMF, as a nonhydrolyzable analog of pTyr, is suitable in cellular studies. The methodology developed in this work will significantly enhance one’s ability to study pTyr-dependent events.

## Results

### Library construction and selection

Based on the analysis of its crystal structure (PDB: 1×8X)^15^, an EcTyrRS mutant library was constructed by randomizing four amino acid residues (Y37, L71, W129, and D182; Supplementary Fig. 1a). These residues likely interact with the carboxylic acid side chain of CMF. The mutant library was achieved by using overlapping PCR and NNK codon (N=A, C, T, or G, K=T or G; 32 variants at nucleotide level) to cover all 20 amino acids at each position. The DNA library fragments were inserted after an ADH1 promoter in a yeast protein expression vector, which also encoded an amber suppressor tRNA derived from *E. coli* tyrosyl-tRNA (*Ec*tRNA_CUA_). By using a yeast homologous recombination cloning method,^16^ a library of 1 × 10^7^ clones was obtained and its quality was verified by DNA sequencing (Supplementary Fig. 1b).^17^

The selection was conducted in *Saccharomyces cerevisiae* by following a previously established procedure.^18-20^ Briefly, the selection was based on the expression of plasmid pGADGAL4-encoded transcriptional activator GAL4, in which codons of Thr44 and Arg110 are replaced with amber codons (Supplementary Fig. 1c). Suppression of the amber codon at both positions were examined in yeast strain MaV203 that contains deletions of the endogenous *GAL4* and *GAL80* genes. Additional chromosomal modifications of MaV203 enable the expression of a genomic copy of *URA3* under the control of a GAL4 responsive promoter. The *URA3* protein complements the uracil auxotrophy of MaV203. The EcTyrRS library was first subjected to a positive selection by culturing cells on synthetic drop-out media plates without uracil to identify EcTyrRS variants that can efficiently aminoacylate the amber suppressor tRNA with CMF. Next, a negative selection was conducted by culturing yeast cells on synthetic drop-out media plates containing 0.1% 5-fluoroorotic acid (5-FOA) without CMF. EcTyrRS variants that can charge the amber suppressor *Ec*tRNA^Tyr^ with any of the endogenous amino acids lead to GAL4-activated expression of URA3, which converts 5-FOA into a cytotoxic product and causes cell death. Growth properties of single colonies survived from the two consecutive selections were examined in synthetic drop-out media either in the presence or in the absence of 1 mM CMF. The ones that only grew in the presence of 1 mM CMF were considered as hits and further characterized.

### Characterization

Four clones (Fig. 1a) were obtained from the selection, from which two unique protein sequences were obtained as CMFRS-1 (Y37H, L71V, and D182G) and CMFRS-2 (Y37H, L71V, W129F, and D182G). Positions Tyr37, Leu71, and Asp182 showed a complete convergence to His, Val, and Gly, respectively, while position Trp129 either maintained the wild-type or changed to Phe. The evolved CMFRSs was evaluated by analyzing the expression of green fluorescent proteins as the reporter in *S. cerevisiae* using a yeast GFP (yeGFP) with an amber mutation at position Asn149. As shown in Fig. 1b, significantly higher (p-value < 0.02) GFP fluorescence was observed in the presence of CMF for both variants. The more prevalent variant, CMFRS-1, led to higher reading. It was designated as CMFRS and chosen for all subsequent experiments.

**Fig. 1.**
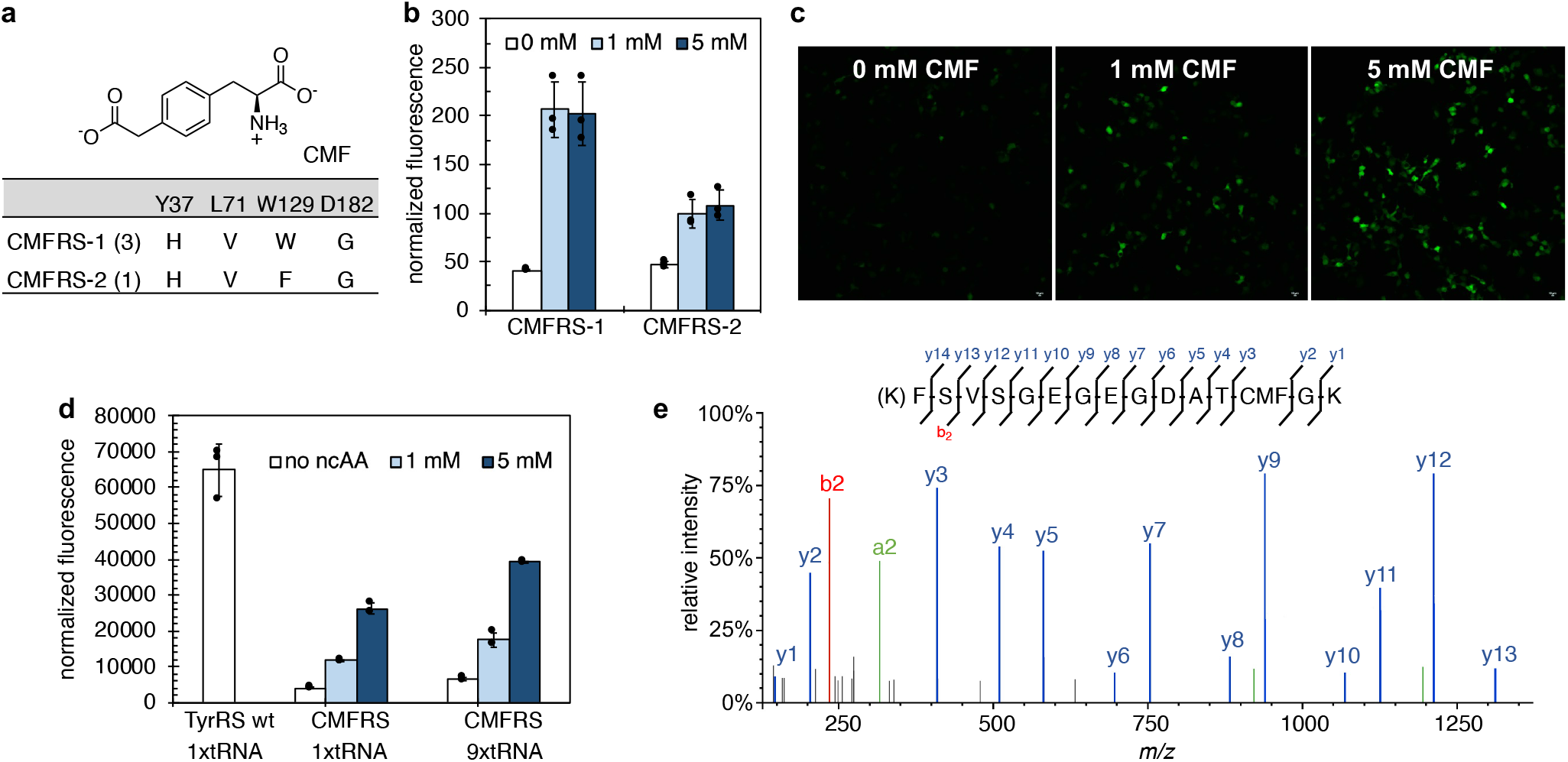
Characterization of the evolved CMFRSs. **a** Mutations in the evolved CMFRS variants. Numbers in parentheses represent the occurrence of the variant in library selection. **b** Fluorescence readings of yeast cells expressing the evolved CMFRSs and an yeGFP mutant that contained an amber mutation at position N149. CMF was included in culture media at 0 mM (open bar), 1 mM (light blue bar), or 5 mM (dark blue bar). Fluorescence intensity was normalized to cell growth. **c** Confocal images of 293T cells expressing the evolved CMFRS and an EGFP mutant that contained an amber mutation at position Tyr40. Scale bars, 10 μm. **d** Flow cytometry analyses of 293T cells expressing the CMFRS or the wild-type EcTyrRS together with the EGFP-Y40TAG mutant. The normalized fluorescence was calculated by multiplying the mean fluorescence intensity by the percentage of fluorescent cells in flow cytometry analyses in the absence (open bar), 1 mM (light blue bar), or 5 mM (dark blue bar) of CMF. **e** LC-MS/MS analysis of the tryptic peptide that contains the position of incorporation. **b** and **d** Data are plotted as the mean ± s.d. from n= 3 independent experiments.

The fidelity and the efficiency of CMFRS was further characterized in 293T cells. For protein expression experiments, the gene that encodes CMFRS was inserted behind a non-regulated CMV promoter on plasmid pCMFRS that contains an amber suppressor tRNA under the control of a human U6 promoter. The selective incorporation of CMF into proteins in 293T cells was then tested by suppression of an amber mutation in a C-terminal His6-tagged EGFP mutant, EGFP-Tyr40TAG, which was encoded on plasmid pEGFP. When cells reached 70% confluency, they were transiently transfected with pCMFRS and pEGFP. After 6 h of incubation at 37 °C in a humidified atmosphere of 5% CO_2_, fresh media was exchanged to include 1 mM or 5 mM CMF and the cells were grown for an additional 24 h. Fluorescence microscopy of the cells revealed that significant EGFP was only expressed in the presence of CMF. Little background expression was observed in the absence of CMF (Fig. 1c). Expression levels of EGFP were further quantified using flow cytometry analysis. The normalized fluorescence was calculated by multiplying the mean fluorescence intensity by the percentage of fluorescent cells. As shown in Fig. 1d, the relative decoding efficiency of CMFRS in the presence of 5 mM CMF is about 40% of the efficiency of the wild-type EcTyrRS.

To further confirm the site-specific incorporation of CMF, peptide sequencing using tandem mass spectrometry (MS/MS) was performed (Fig. 1e). To this end, the EGFP-Tyr40TAG mutant was expressed in 293T cells in the presence of 5 mM CMF, purified by affinity chromatography, and further separated from impurities by SDS-PAGE (Supplementary Fig. 2a). Following trypsin digest, the peptide fragment that contained the position of incorporation was analyzed. Fragmentation data provided clear evidence that CMF was incorporated at position 40 of EGFP (Fig. 1e). Finally, the fidelity of incorporation by the evolved CMFRS was quantified based on the relative abundance of the modified and unmodified peptides (Supplementary Fig. 2b). The precursor ions for peptide containing CMF (773.3395 m/z, [M+2H^+^]) and peptide containing Tyr (752.3345 m/z, ([M+2H^+^]) were extracted at 5 ppm mass accuracy. Comparison of integrated peak areas showed that the CMF-containing peptide was detected at a 98% rate.

### Optimization of the decoding system

It was reported that the expression level of the tRNA is the limiting factor of decoding efficiency in mammalian cells.^21,22^ To further increase the CMF incorporation, a series of pCMFRS plasmid variants with different copies of tRNA were constructed. These plasmids were co-transfected with plasmid pEGFP 2TAG that contains the EGFP-40TAG-150TAG for the evaluation of multi-site CMF incorporation in 293T cells. Fluorescence of EGFP was observed in the presence of both 1 mM and 5 mM CMF. Significantly higher (*p* < 0.01) level of expression was observed at 5 mM (Supplementary Fig. 2c). Furthermore, the EGFP expression increases with increasing copies of tRNA (Supplementary Fig. 2c). The CMFRS/9xtRNA plasmid was also examined together with plasmid pEGFP by flow cytometry. A decoding efficiency of around 60% of that of the wild-type EcTyrRS was observed (Fig. 1d). This construct was used in subsequent studies.

### DNA binding study

We examined if functional replacement of pTyr with CMF can be realized in the human signal transducer and activator of transcription-1 (STAT1) protein. Phosphorylation at the Tyr701 position of STAT1 leads to its formation of a homodimer and binding to specific DNA sequences (e.g.,γ-IFN-activated sequence) following nuclear translocation.^23^ In this experiment, the DNA binding capability of the C-terminal fragment of STAT1 mutant containing CMF at position 701 (132-712, c-STAT1-701CMF) was studied *in vitro*. The c-STAT1-701CMF was expressed and purified from 293T cells (Supplementary Fig. 3a and 3b). The site specific incorporation of CMF at position 701 was verified by MS/MS analysis (Supplementary Fig. 3c). The interaction between c-STAT1-CMF701 mutant and a short M67 DNA fragment (Fig. 2) was examined by fluorescence-based electrophoretic mobility shift assay (EMSA). As shown in Fig. 2, gel shifting of DNA caused by protein binding was observed when c-STAT1-CMF701 concentration reached 240 nM. Meanwhile, the control experiment using the unphosphorylated c-terminal fragment of the wild-type STAT1 protein (c-STAT1-701Tyr) did not lead to gel shifting in the presence of the same concentration of M67 DNA fragment. The findings confirmed that CMF can functionally replace pTyr at the 701 position in STAT1 protein that was expressed in mammalian cells.

**Fig. 2.**
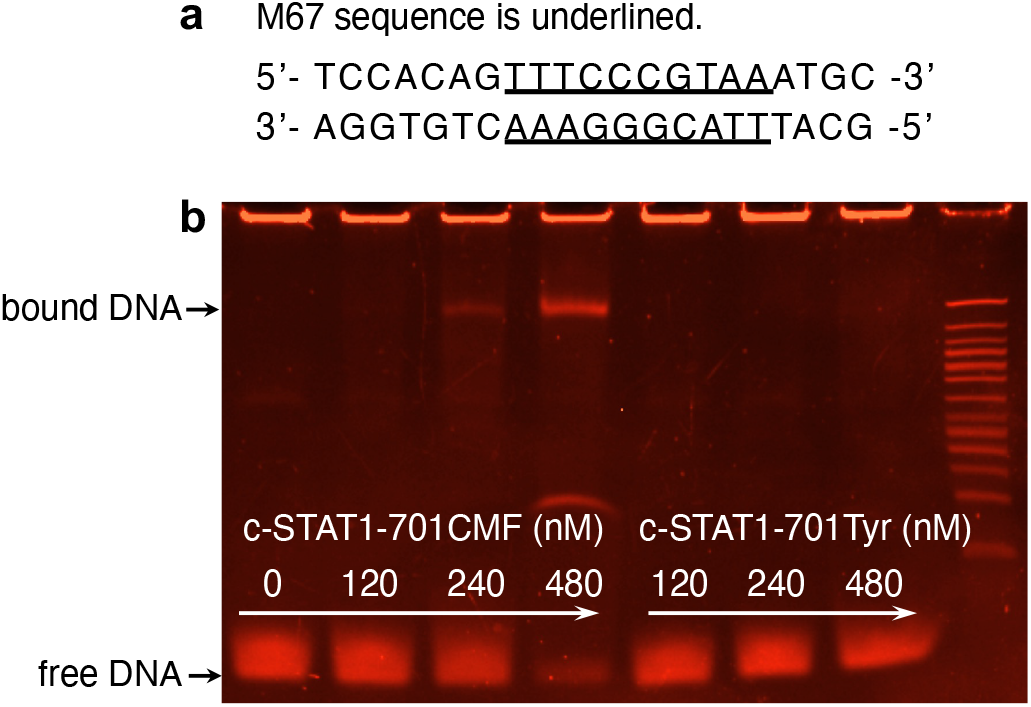
Interaction between STAT1 and its DNA substrate. **a** Sequence of the 21-bp DNA oligonucleotide duplex used in this study. The M67 site, which is recognized by STAT1 dimer following phosphorylation at Try701, is underlined. **b** Fluorescence-based EMSA assay. The DNA concentration was fixed at 30 nm. The protein concentrations varied from 0 to 480 nM. Free and bound (DNA-protein complex) DNAs were visualized by SYBR® Gold EMSA nucleic acid gel stain.

### Antibody study

To further demonstrate that the STAT1-701CMF mutant functions in a similar manner to naturally phosphorylated STAT1, we examined the behavior of STAT1-701CMF with phospho-STAT1 (pTyr701) antibody. We first conducted western blot experiments using the total lysate of 293T cells that expressed variants of the STAT1-EGFP fusion protein. As shown in Fig. 3a (lane 4), a protein band was clearly detected by the phospho-STAT1 (pTyr701) antibody when STAT1-701TAG-EGFP was expressed in the presence of CMF. The detected protein has a similar size to the phosphorylated STAT1-EGFP fusion protein, which was observed in interferon-γ (IFN-γ)-induced, STAT1-EGFP-expressing cells (lane 2, Fig. 3a). On the other hand, the pTyr701 antibody did not detect such protein band when STAT1-701TAG-EGFP was expressed in the absence of CMF (lane 3, Fig. 3a). Similar result was observed in the negative control, cells expressing STAT1-EGFP without the induction of IFN-γ (lane 1, Fig. 3a). Further western blot experiments with anti-STAT1 antibody (independent of phosphorylation state) verified the expression and the correct size of STAT1-EGFP variants (lanes 5 to 8, Fig. 3a). A low level of expression was detected in the lysate of STAT1-701TAG-EGFP cells without exogenously added CMF (lane 7, Fig. 3a). These observations confirmed that STAT1-701CMF can be recognized by phospho-STAT1 (pTyr701) antibody.

**Fig. 3.**
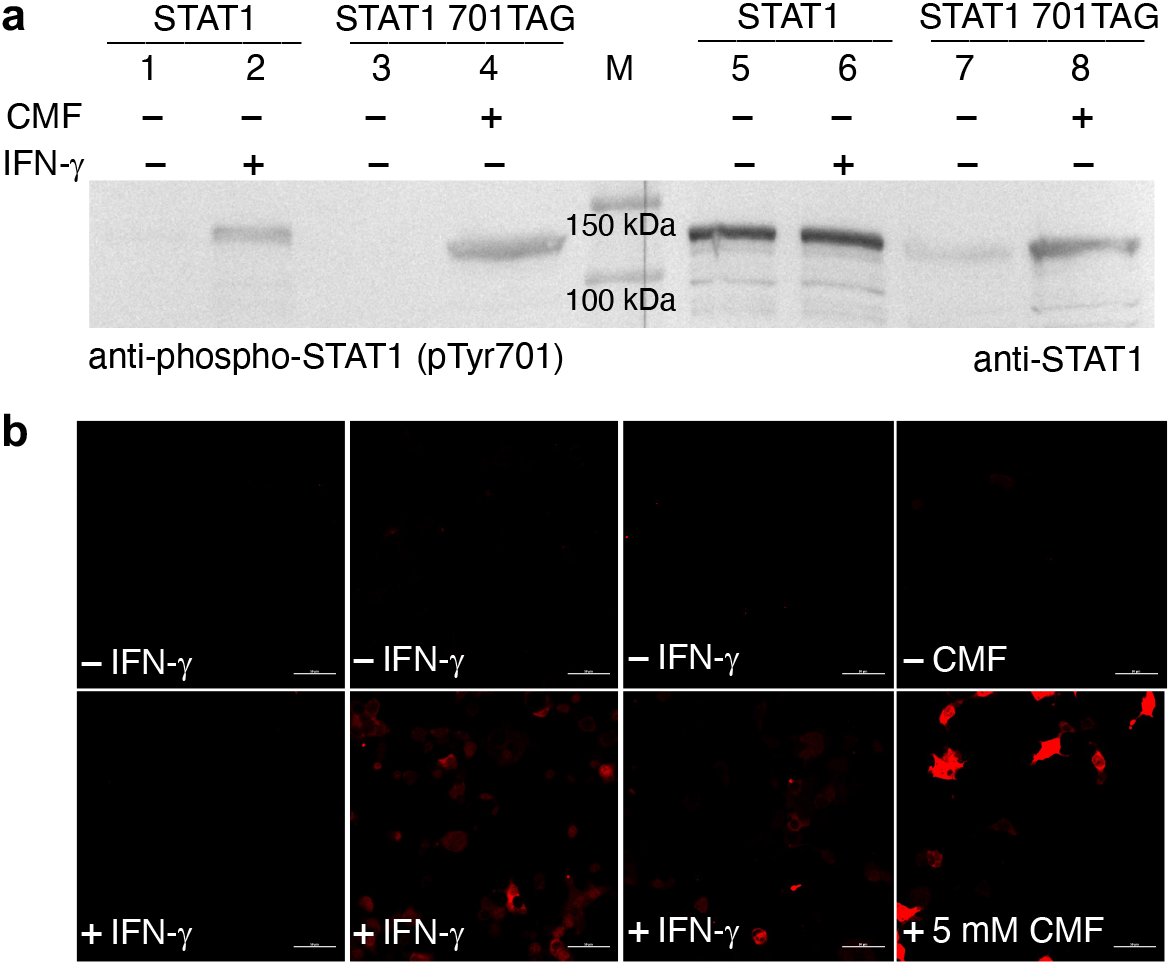
Detection of STAT1 by phospho-STAT1 (pTyr701) antibody. **a** Western blot of cell lysates expressing STAT1 variants. Lanes 1 and 5, wild-type STAT1-EGFP without IFN-γ induction; lanes 2 and 6, wild-type STAT1-EGFP with IFN-γ induction; lanes 3 and 7, STAT1-701TAG-EGFP without CMF; lanes 4 and 8, STAT1-701TAG-EGFP with CMF. Lane M is the molecular weight marker. The calculated molecular weight of STAT1-EGFP variants was 115.9 kDa. Lanes 1-4 were detected by phospho-STAT1 (pTyr701) antibody. Lanes 5-8 were detected by anti-STAT1 antibody. **b** Images of cells expressing STAT1 variants. Left panel 293T-ΔSTAT1 cells; mid-left panel, 293T-ΔSTAT1 cells expressing wild-type STAT1; mid-right panel, 293T-ΔSTAT1 cells expressing STAT1-701TAG and the wild-type EcTyrRS; right panel, 293T-ΔSTAT1 cells expressing STAT1-701TAG and the CMFRS. Immunofluorescence images were obtained by the treatment of permeabilized cells with phospho-STAT1 (pTyr701) antibody followed by anti-rabbit secondary antibody-Alex594 conjugate. Scale bars, 50 μm.

Next, we conducted cell-based immunofluorescence assay. In order to remove background signals from endogenously expressed STAT1, we first generated a STAT1 knockout cell line, 293T-ΔSTAT1, using CRISPR/Cas method (see details in Methods).^24^ Cells were treated and cultivated under indicated conditions (Fig. 3b), then fixed and permeabilized. The phospho-STAT1 (pTyr701) antibody was added as the primary antibody, followed by an Alexa Fluor 594-labeled secondary antibody. Cells were then visualized under the confocal microscope. As shown in the left panel of Fig. 3b, no signal was detected with 293T-ΔSTAT1 cells either in the absence or in the presence of IFN-γ induction. This result confirmed the successful deletion of the STAT1-encoding gene in this cell line. Following transfection with pSTAT1, fluorescence was detected only in cells that were induced with IFN-γ (mid-left panel in Fig. 3b), which verified a functional signaling pathway in the cell line. Meanwhile, strong signal was observed when STAT1-701TAG was expressed together with CMFRS in the presence of CMF (lower right panel in Fig. 3b) and no fluorescent cells were detected in the absence of CMF (upper right panel in Fig. 3b). This observation is in line with control cells that expressed STAT1-701TAG together with the wild-type EcTyrRS, for which fluorescence was only detected in IFN-γ-induced cells (mid-right panel in Fig. 3b). Results of the immunofluorescence assay therefore confirmed that STAT1-701CMF functions similarly to STAT1-701pTyr in live mammalian cells.

## Discussion

This work represents the first genetic encoding of a nonhydrolyzable pTyr analog in yeast and mammalian cells. While the evolved CMFRS mutants did display background activity, mis-incorporation of Tyr was low (< 2%) in the presence of 5 mM CMF. The evolved CMFRS shares two common mutations, L71V and D182G, with a sTyrRS that we reported previously for the encoding of sulfotyrosine (sTyr) in mammalian cells.^17^ Analysis of the crystal structure of sTyrRS in complex with its substrate (PDB ID: 6WN2) have shown that both mutations expand the active site in order to accommodate the large substituent at the *para*-position of Tyr. In addition, the D182G mutation eliminates the electrostatic repulsion between the negatively charged Asp and the substituent on Tyr.^17^ These arguments stay true to the substrate binding by the CMFRS mutants. A model of CMFRS was generated by introducing the additional mutations into the structure of sTyrRS. Docking of CMF into the active site shows that the Y37H mutation, which is unique to both CMFRS mutants, leads to hydrogen bond formation between the carboxylate group of CMF and the nitrogen on the imidazole side chain (Supplementary Fig. 1d). The interaction is apparently essential for the substrate recognition by CMFRSs.

In a proof-of-concept study using STAT1 as the model protein, we for the first time provided evidence to show that CMF could serve as a functional analog of pTyr in mammalian cells. Results of the *in vitro* EMSA assay using proteins expressed in 293T cells verified that CMF incorporated at the position 701 of STAT1 mimics the function of pTyr by triggering protein dimerization and its subsequent binding to the DNA substrate. Western blot experiments using cell lysate further demonstrated that the STAT1-701CMF could be recognized by phospho-STAT1 (pTyr701) antibody, which indicates a strong structural and electrostatic similarity to STAT1-701pTyr. The immunofluorescence study using live cells also validated the recognition by pTyr701-specific antibody. Furthermore, stronger signal was observed in individual cells that expressed STAT1-701CMF in comparison to cells expressing wild-type STAT1 under IFN-γ induction. Since the phosphorylation state of Tyr701 in STAT1 is dynamically controlled by the kinase and phosphatase activities, only a sub-population of cellular STAT1 protein is at the pTyr701 state.^23^ Meanwhile, as a nonhydrolyzable pTyr analog, STAT1 protein with CMF at 701 position sustains its chemical stability and the ability to bind the antibody independent of the endogenous regulatory mechanism. Overall, this work makes the efficient expression of proteins containing a nonhydrolyzable pTyr analog in yeast and mammalian cells possible, thereby enabling structure-function studies as well as the practical therapeutic application of phosphoproteins. We expect our work not only to facilitate fundamental investigations, but also to accelerate biomedical applications that target protein tyrosine phosphorylation.

### Significance

Protein tyrosine phosphorylation plays essential roles in the regulation of cellular activities. The dynamic nature of this posttranslational modification arises from activities of protein kinases and phosphatases, which complicates the study of a particular tyrosine phosphorylation event. In this report, we established, for the first time, a method for the genetic incorporation of a nonhydrolyzable analog of phosphotyrosine, *p*-carboxymethyl-L-phenylalanine (CMF), in live mammalian cells. Our protein engineering efforts led to a mutant *E. coli* tyrosyl tRNA synthetase (CMFRS) that showed excellent activity and specificity for CMF. Using the human signal transducer and activator of transcription-1 (STAT1) as a model protein, we demonstrated that the incorporation of CMF at position 701, which is a natural tyrosine phosphorylation position that triggers the activation of STAT1, led to proteins with biological activities that engaged in interactions with the target DNA sequence and phosphotyrosine-specific antibody. The method developed in this report has broad applicability for live cell studies or the production of therapeutics where tyrosine phosphorylation is critical to the biological functions.

## Acknowledgments

The authors thank Dr. You Zhou and Ms. Terri Fangman (Morrison Microscopy Core Research Facility at University of Nebraska-Lincoln) for assistance in confocal microscope analysis, Dr. Dirk Anderson (Flow Cytometry Service Center at University of Nebraska-Lincoln) for assistance in flow cytometry analysis, and Drs Sophie Alvarez and Mike Naldrett (Proteomics and Metabolomics Facility) for mass spectrometry analysis. This work was supported by National Science Foundation (grant CBET 1805528 to W.N. and J.G.), National Science Foundation (grant MCB 1553041 to J.G.), National Institute of Health (grant 1R01GM138623 to J.G. and W.N.).

## Author Contributions

W.N. and J.G. designed the study. X.H. constructed plasmids and the EcTyrRS library and performed selection in yeast, purified protein and conducted EMSA, western blot, and immunofluorescence assay. B.M. synthesized CMF. Y.C. conducted confocal microscopy and flow cytometry studies. X.H., J.G., and W.N. analyzed the data and wrote the manuscript.

## Declaration of interests

The authors declare no competing interests.

## STAR Methods

### Resource availability

#### Lead contact

Further information and requests for rsources and reagents should be directed to the Lead Contact: Wei Niu and Jiantao Guo

#### Materials availability

Requests for materials should be addressed to W.N. and J.G.

#### Data availability

The article includes all datasets generated or analyzed during this study.

#### Experimental model and subject details

Human embryonic kidney cells (HEK293T, ATCC: CRL-3216) were used for this study. The cells were maintained in Dulbecco’s modified Eagle’s medium (DMEM) supplemented with 10% fetal bovine serum (FBS) at 37 °C in a humidified atmosphere of 5% CO_2_ (v/v).

### Method details

#### Reagents and media

All commercial chemicals are of reagent grade or higher. All solutions were prepared in deionized water that was further treated by Barnstead Nanopure® ultrapure water purification system (Thermo Fisher Scientific). Preparation of LB medium and YPD medium followed reported recipe. Yeast selection media contained yeast nitrogen base without amino acids (Difco^™^, 6.7 g/L), glucose (10 g/L), and appropriate drop out (DO) supplements (Clonetech). The pH values of all media were adjusted to 7.0. Agar plates were prepared by the addition of Difco agar (15 g/L) to the liquid medium. Antibiotics were added where appropriate to following final concentrations: kanamycin (50 mg/L), ampicillin (100 mg/L). Solutions of antibiotics were filtered through 0.22 μm sterile membrane filters. CMF was synthesized by following previously published procedure with modifications.^10^ (Supplementary Information).

#### Cell culture and transfection

HEK293T cells and 293T-ΔSTAT1 were maintained in DMEM media supplemented with 10% FBS at 37 °C in a humidified atmosphere of 5% CO_2_ (v/v). Transfection was conducted at 70-80% cell confluency using Lipofectamine 2000 (Life Technologies) according to the manufacturer’s protocol.

#### Strains and plasmids

*E. coli* GeneHogs^®^ (Thermo Fisher Scientific) was used for routine cloning and plasmid propagation unless otherwise mentioned. *E. coli* NEB^®^ Stable (New England BioLabs) was used for cloning and propagation of unstable plasmids. Plasmid construction was performed using T4 DNA ligase or Gibson Assembly method.^25^ *S. cerevisiae* strain MaV203 (MATα; leu2-3,112; trp1-109; his3Δ200; ade2-101; cyh2R; cyh1R; gal4Δ; gal80 Δ; GAL1::lacZ; HIS3UASGAL1::HIS3@LYS2; SPAL10UASGAL1::URA3) (Thermo Fisher Scientific) was used for yeast library construction and selection. Standard protocols were followed for the purification and analysis of plasmid DNA.^26^ PCR amplifications were carried out using KOD Hot Start DNA polymerase (Millipore Sigma) by following the manufacturer’s protocol. Other molecular cloning reagents were purchased from New England BioLabs. Primer synthesis service was provided by Millipore Sigma, and DNA sequencing service was provided by Eurofins MWG Operon. Primers used in this study are listed in Supplementary Table 1. Plasmids used and constructed in this study are listed in Supplementary Table 2. All plasmids are available from corresponding authors upon reasonable request.

#### 293T-ΔSTAT1 cell

A STAT1-knockout cell line was generated from 293T using CRISPR/Cas9 method. The gRNA sequence^27^ (5’-TCATGACCTCCTGTCACAGC-3’) was inserted into BbsI-linearized plasmid pSpCas9(BB)-2A-GFP (PX458, Addgene 48138). The resulting plasmid PX458-STAT1 gRNA was transfected into 293T cells. 24 h post transfection, cells were detached and filtered through a 35 μm cell strainer and single cells with GFP expression were sorted into a 96-well plate using BD FACSAria II. Colonies developed from single cells were expanded and maintained. A pair of primers flanking gRNA binding site were used to amplify PCR products for DNA sequencing in order to identify chromosomal editing events. The selected 293T STAT1-knockout cell line had a 365-bp deletion on both chromosomes, which covered 99 bp of exon and 266 bp of intron. The phenotype of the cell line was confirmed by western blot.

#### Plasmid pCMFRS

This 7.5-kb pcDNA3.1-derived plasmid contains one copy of the amber suppressor *B. subtilis* tyrosyl-tRNA under the control of a human U6 promoter and a CMFRS-encoding gene behind a CMV promoter. Fragment encoding the CMFRS-1 was amplified from the isolated pEcTyrRS-CMFRS1 and inserted into a 6.2-kb vector that was obtained after treating pcDNA3.1-AzFRS with HindIII and ApaI.

#### Plasmids pSTAT1-EGFP and derivatives

The 9.2-kb pSTAT1-EGFP plasmid is derived from pcDNA3.1 by replacing the AzFRS-encoding gene in pcDNA3.1-AzFRS with DNA fragment of human STAT1 and EGFP fusion protein. STAT1 and EGFP genes were connected by a short DNA sequence that encodes a GSGSGSAAA linker. To make pSTAT1 Y701TAG-EGFP, TAG were introduced at the Y701 position of the STAT1 gene in pSTAT1-EGFP by mutagenic PCR.

#### Plasmids pSTAT1 and pSTAT1 Y701TAG

The 8.4-kb pSTAT1 and pSTAT1-Y701TAG plasmids were constructed by inserting the PCR products of STAT1 and STAT1 Y701TAG into a 6.2-kb vector that was derived from HindIII and ApaI treatment of pSTAT1-EGFP, respectively.

#### Plasmids pSTAT1 (132-712) and pSTAT1 (132-712) Y701TAG

The 8.0-kb pSTAT1 (132-712) and pSTAT1 (132-712) Y701TAG plasmids are derived from pSTAT1-EGFP. DNA fragments encoding residues 132-712 of STAT1 and STAT1 701TAG were amplified, purified, and assembled into a 6.2-kb DNA fragment from the digestion of pSTAT1-EGFP by HindIII and XhoI, respectively.

#### Plasmid pSpCas9(BB)-2A-GFP-STAT1 gRNA

The 9.3-kb pSpCas9(BB)-2A-GFP-STAT1 gRNA plasmid is derived from pSpCas9(BB)-2A-GFP (PX458, Addgene). Primers STAT1 gRNA-F and STAT1 gRNA-R were annealed and phosphorylated following standard protocol.^24^ The mixture was ligated into the BbsI-linearized plasmid pSpCas9(BB)-2A-GFP. *E. coli* NEB^®^ Stable was used for efficient transformation and stable propagation.

#### Plasmids pCMFRS with multiple copies of Bs tRNA

The 8.2-kb plasmid pCMFRS-3xtRNA contains three copies of the *B. subtilis* amber suppressor tyrosyl-tRNA under the control of a human U6 promoter and a human H1 promoter. Promoter H1 was amplified from 293T genome and assembled with Bs tRNA as a H1-Bs tRNA gene cassette by overlapping PCR. The U6-Bs tRNA cassette was amplified from pCMFRS. The two cassettes were inserted into BstZ17I-digested pCMFRS to obtain pCMFRS-3xtRNA. The 8.9-kb pCMFRS-5xtRNA was constructed by ligating the H1-Bs tRNA-U6-Bs tRNA fragment, which was released from pCMFRS-3xtRNA by NheI/XbaI, into the XbaI-treated pCMFRS 3xtRNA. The same cloning strategy for pCMFRS-5xtRNA was applied to the construction of plasmids pCMFRS-7xtRNA and pCMFRS-9xtRNA using pCMFRS-5xtRNA and pCMFRS-7xtRNA as the vector, respectively. *E. coli* NEB^®^ Stable was used as the cloning host.

#### Library construction and selection

The EcTyrRS library was constructed previously in MaV203/pGADGAL4 by randomizing four residues (Y37, L71, W129, and D182) using the NNK codon (N = A, C, T, or G, K = T or G).^17,28^ A library size of 1 × 10^7^ diversity was obtained and maintained in SD-Leu-Trp media. Positive selection was conducted by culturing cells on the SD-Leu-Trp-Ura plates with 1 mM CMF to identify EcTyrRS variants that can efficiently aminoacylate the amber suppressor tRNA with CMF. In the negative selection, cells were plated on SD-Leu-Trp media plates containing 5-fluoroorotic acid (5-FOA) at 0.1%. EcTyrRS variants that can charge the amber suppressor *Ec*tRNA_CUA_ with any one of the 20 natural amino acids led to GAL4-activated expression of URA3, which converts 5-FOA into a cytotoxic product and leads to cell death. Single colonies picked from the selection plates were resuspended in 100 μL of SD-Leu-Trp-Ura media. A 2 μL cell resuspension was applied to SD-Leu-Trp-Ura, and SD-Leu-Trp-Ura plates with 1 mM CMF to further verify the phenotypes. Plasmids isolated from potential candidates were propagated in *E. coli* GeneHogs for DNA sequencing.

#### Fluorescence analysis of yeast culture

MaV203 transformed with plasmids pyeGFP-N149TAG and pEcTyrRS-mutant was cultured in SD-Leu-Trp media without or with CMF (1 mM or 5 mM) at 30 °C for 12 h. Cells were then collected, washed, and resuspended in an equal volume of PBS (pH 7.4). Cell density was determined by measuring OD_600nm_. The fluorescence of yeGFP was monitored at Ex = 485 nm and Em = 528 nm. Values of fluorescence intensity were normalized to the OD measurement. Reported data are the average of three measurements with standard deviations. A Biotek Synergy HTX plate reader was used in absorbance, and fluorescence measurements.

#### Confocal microscopy

293T cells (1 × 10^5^) seeded in a single well of a 24-well plate were grown for 16-24 h, then transfected with plasmids pCMFRS (0.8 μg) and pEGFP (0.8 μg) in 2 μL of Lipofectamine. Transfected cells were cultured for an additional 24 h in 0.5 mL of media without or with CMF (1 mM or 5 mM). Cells were then washed with 0.5 mL of warm DMEM base medium, fixed with 4% paraformaldehyde (w/v) for 15 min. Following removal of the fixation reagent by washing with 3 × 0.5 mL DPBS, cells were visualized by an Inverted (Olympus IX 81) confocal microscope.

#### Flow cytometry analysis

For the quantification of amber suppression efficiency by CMFRS, transfected 293T cells from a single well of a 24-well plate were detached with 0.3 mL of 0.05% Trypsin/EDTA, washed with 0.5 mL DPBS, and collected by centrifugation at 300 x *g* for 5 min. Collected cells were fixed in 0.5 mL 4% paraformaldehyde (w/v) for 15 min at room temperature. Following removal of the fixation reagent by centrifugation, cells were washed twice and resuspended in 0.5 mL DPBS and kept on ice until analysis. Fluorescence of cells were measured using a Beckman Coulter CytoFLEX flow cytometer. A total of 30,000 cells were analyzed for each sample. Data were analyzed using FlowJo. Reported data are the average measurement of three samples with standard deviations.

#### Mass spectrometry analysis

Proteins EGFP-40CMFand c-STAT1-701CMF were overexpressed in 293T cells transfected with pCMFRS together with pEGFP or pSTAT1 (132-712) Y701TAG. Cells were lysed by sonication in protein binding buffer (25 mM Tris-HCl, 200 mM NaCl, 10 mM imidazole, 1 x protease inhibitor cocktail, pH 7.5). Following centrifugation at 21,000 x *g* for 30 min at 4 °C, tagged protein was first purified using Ni Sepharose 6 Fast Flow resin, then resolved from impurity by SDS-PAGE. Bands corresponding to target proteins were excised, washed, and treated with trypsin overnight at 37 °C. Tryptic peptides were extracted from the gel pieces, dried down, and re-dissolved in 25 μL of an aqueous solution of acetonitrile (2.5%) and formic acid (0.1%).

Each digest was run by nano LC-MS/MS using a 1-h gradient on a 0.075 mm x 250 mm C18 column feeding into a Q-Exactive HF mass spectrometer. All MS/MS samples were analyzed using Mascot. Mascot was set up to search a database that was customized with provided protein sequence. Mascot search has a fragment ion mass tolerance of 0.060 Da and a parent ion tolerance of 10.0 ppm. Deamidation of asparagine and glutamine, oxidation of methionine, and CMF incorporation were specified in Mascot as variable modifications. Scaffold was used to validate MS/MS based peptide and protein identifications. Peptide identifications were accepted if they could be established at greater than 99.0% probability by the Peptide Prophet algorithm^29^ with Scaffold delta-mass correction. Protein identifications were accepted if they could be established at greater than 99.0% probability and contained at least one identified peptide. Protein probabilities were assigned by the Protein Prophet algorithm.^30^

#### Western blot

Harvested 293T cells were lysed with RIPA buffer (Thermo Scientific) containing Halt^™^ protease and phosphatase inhibitor cocktail (Thermo Scientific). Following centrifugation, the supernatant was saved as cell extracts. Protein concentrations were then determined using the Quick Start^™^ Bradford protein assay kit (Bio-Rad Laboratories). Proteins were separated by SDS-PAGE, then transferred onto a nitrocellulose membrane using mini trans-blot cell (Bio-Rad Laboratories). The membrane was blocked with 3% BSA at 4 °C overnight, incubated with primary antibodies at 4 °C for 8 hours, and incubated with secondary antibodies at room temperature for 1 h. Blots were developed using Opti-4CN detection kit (Bio-Rad Laboratories) and imaged using Gel Doc XR+ system (BioRad Laboratories). Primary antibodies used in this study included rabbit anti-phospho-STAT1 (pTyr701) (D4A7) (1:1000, Cell Signaling Technology), mouse anti-STAT1 antibody (C-111) (1:500, Santa Cruz Biotechnology), and mouse anti-histidine tag antibody (clone AD1.1.10) (1:3000, Bio-Rad Laboratories). Secondary antibodies used in this study included HRP conjugated goat anti-mouse IgG (1:2000, Bio-Rad Laboratories) and HRP-linked goat anti-rabbit IgG (1:2000, Cell Signaling Technology).

#### Electrophoretic mobility shift assay (EMSA)

Proteins c-STAT1 and c-STAT1-701CMF were overexpressed and purified from 293T cells transfected with pSTAT1 (132-712) or pCMFRS together with pSTAT1 (132-712) Y701TAG. The same protocol of protein purification for MS analysis was followed. Imidazole was removed from purified protein by dialysis against a storage buffer (25 mM Tris-HCl, 150 mM NaCl, 5% glycerol, 2 mM DTT, pH 7.5). Concentrations of purified c-STAT1 variants were determined by western blot. Calibration curve was constructed using c-STAT1 purified from *E. coli* host^31^ with known concentrations.

EMSA followed a previously reported protocol.^31^ PAGE-purified M67-1 and M67-2 primers were dissolved in annealing buffer (5 mM Tris-HCl, 50 mM KCl, 10 mM MgCl_2_, pH 8.0) to a final concentration of 4 μM each. The mixture was heated to 94 °C, then slowly cooled to room temperature to form DNA duplex. In EMSA assay, 30 nM M67 was mixed with indicated concentrations of c-STAT1 variants in reaction buffer (20 mM HEPES, 4% Ficoll, 40 mM KCl, 10 mM CaCl_2_, 10 mM MgCl_2_, 1 mM DTT, 0.2 mg/mL BSA, pH 8.0). The reaction mixture was incubated at room temperature for 30 min, then resolved on a pre-run 4-20% Novex TBE gel for 40 min at 4 °C. The gel was stained with SYBR^™^ Gold and visualized using Gel Doc^™^ XR+ imaging system.

#### Immunofluorescence assay

The 293T-ΔSTAT1 cell line was used for immunofluorescence assay. Cells were seeded in poly-lysine coated, clear bottom, black wall 96-well plate (Greiner Bio-One). For IFN-γ induction, a final concentration of 100 ng/μL of IFN-γ (Cell Signaling Technology) was added into cell culture 30 min prior to fixation using 4% formaldehyde. Fixed cells were permeabilized with ice-cold 100% methanol at −20 °C for 10 min, incubated with blocking buffer (5% BSA and 0.1% Triton X-100 in DPBS) at room temperature for 1 h, incubated with anti-phospho-STAT1 (pTyr701) antibody (1:50 dilution, 2% BSA and 0.1% Triton X-100 in DPBS) at 4 °C for 12-16 h, then incubated with the Alexa Fluor^®^ 594 conjugated-anti-rabbit IgG (1:500 dilution, 2% BSA and 0.1% Triton™ X-100 in DPBS) at room temperature for 2 h. Buffer of 0.1% Triton X-100 in DPBS (100 μL) was used for washing between each steps and after the incubation with the fluorophore-conjugated secondary antibody. Cells were visualized by an inverted (Olympus IX 81) confocal microscope.

### Statistical analysis

Standard deviations and Student’s t test were performed using Microsoft Excel 365 software.

## Additional information

Supplementary information is available for this paper.

## Notes

### Competing Interest Statement

The authors have declared no competing interest.

